# Deep Learning-Based Approaches for Decoding Motor Intent from Peripheral Nerve Signals

**DOI:** 10.1101/2021.02.18.431483

**Authors:** Diu Khue Luu, Anh Tuan Nguyen, Ming Jiang, Jian Xu, Markus W. Drealan, Jonathan Cheng, Edward W. Keefer, Qi Zhao, Zhi Yang

**Affiliations:** Biomedical Engineering, University of Minnesota, Minneapolis, MN, USA; Computer Science and Engineering, University of Minnesota, Minneapolis, MN, USA; Plastic Surgery, University of Texas Southwestern Medical Center, Dallas, TX, USA; Nerves Incorporated, Dallas, TX, USA; Fasikl Incorporated, Minneapolis, MN, USA

**Keywords:** convolutional neural network, deep learning, feature extraction, motor decoding, neuroprosthesis, neural decoder, peripheral nerve interface, recurrent neural network

## Abstract

The ultimate goal of an upper-limb neuroprosthesis is to achieve dexterous and intuitive control of individual fingers. Previous literature shows that deep learning (DL) is an effective tool to decode the motor intent from neural signals obtained from different parts of the nervous system. However, it still requires complicated deep neural networks that are inefficient and not feasible to work in real-time. Here we investigate different approaches to enhance the efficiency of the DL-based motor decoding paradigm. First, a comprehensive collection of feature extraction techniques is applied to reduce the input data dimensionality. Next, we investigate two different strategies for deploying DL models: a one-step (1S) approach when big input data are available and a two-step (2S) when input data are limited. With the 1S approach, a single regression stage predicts the trajectories of all fingers. With the 2S approach, a classification stage identifies the fingers in motion, followed by a regression stage that predicts those active digits’ trajectories. The addition of feature extraction substantially lowers the motor decoder’s complexity, making it feasible for translation to a real-time paradigm. The 1S approach using a recurrent neural network (RNN) generally gives better prediction results than all the ML algorithms with mean squared error (MSE) ranges from 10^−3^ to 10^−4^ for all finger while variance accounted for (VAF) scores are above 0.8 for the most degree of freedom (DOF). This result reaffirms that DL is more advantageous than classic ML methods for handling a large dataset. However, when training on a smaller input data set as in the 2S approach, ML techniques offers a simpler implementation while ensuring comparably good decoding outcome to the DL ones. In the classification step, either machine-learning (ML) or DL models achieve the accuracy and F1 score of 0.99. Thanks to the classification step, in the regression step, both types of models result in comparable MSE and VAF scores as those of the 1S approach. Our study outlines the trade-offs to inform the implementation of real-time, low-latency, and high accuracy DL-based motor decoder for clinical applications.

## 1 Introduction

Upper-limb amputation affects the quality of life and well-being of millions of people in the United States, with hundreds of thousand new cases annually (Ziegler-Graham, 2008). Neuroprosthetic systems promise the ultimate solution by developing human-machine interfaces (HMI) that could allow amputees to control robotic limbs using their thoughts (Schultz, 2011; Cordella, 2016). It is achieved by decoding the subject’s motor intent with neural data acquired from different parts of the nervous system. Proven approaches include surface electromyogram (EMG) (Sebelius, 2005; Jiang, 2012; Fougner, 2012; Amsuss, 2013; Zuleta, 2019), electroencephalogram (EEG) (Hu, 2015; Zeng, 2015; Sakhavi, 2018; Kwon, 2019), cortical recordings (Mollazadeh, 2011; Hochberg, 2012; Irwin, 2017), and peripheral nerve recordings (Micera, 2011; Davis, 2016; Vu, 2017; Wendelken, 2017; Zhang, 2017; Nguyen & Xu, 2020).

However, implementing an effective HMI for neuroprostheses remains a challenging task. The decoder should be able to predict the subject’s motor intents accurately and satisfy certain criteria to make it practical and useful in daily lives. Some criteria include *dexterity*, i.e. controlling multiple degrees-of-freedom (DOF) such as individual fingers; *intuitiveness*, i.e. reflecting the true motor intent in mind, and *real-time*, i.e. having minimal latency from thoughts to movements.

In recent years, deep learning (DL) techniques have emerged as strong candidates to overcome this challenge thanks to their ability to process and analyze biological big data (Mahmud, 2018). Our previous work (Nguyen & Xu, 2020) shows that neural decoders based on the convolutional neural network (CNN) and recurrent neural network (RNN) architecture outperform other “classic” machine learning counterparts in decoding motor intents from peripheral nerve data obtained with an implantable bioelectric neural interface. The DL-based motor decoders can regress the intended motion of fifteen degrees-of-freedom (DOF) simultaneously, including flexion/extension and abduction/adduction of individual fingers state-of-the-art performance metrics, thus complying with the dexterity and intuitiveness criteria.

Here we build upon the foundation of (Nguyen & Xu, 2020) by exploring different strategies to optimize the motor decoding paradigm’s efficiency. The aim is to lower the neural decoder’s computational complexity while retaining high accuracy predictions to make it feasible to translate the motor decoding paradigm to real-time operation suitable for clinical applications, especially when deploying in a portable platform.

First, we utilize feature extraction to reduce data dimensionality. By examining the data spectrogram, we learn that most of the signals’ power concentrates in the frequency band 25-600 Hz (Nguyen & Xu, 2020). Many feature extraction techniques (Zardoshti-Kermani, 1995; Phinyomark, 2009; Rafiee, 2011; Phinyomark, 2012) have been developed to handle signals with similar characteristics. Feature extraction aims not only to amplify the crucial information, lessen the noise but also to substantially reduce the data dimensionality before feeding them to DL models. This could simultaneously enhance the prediction accuracy and lower the DL models’ complexity. Here we focus on a comprehensive list of fourteen features that consistently appear in the field of neuroprosthesis.

Second, we explore two different strategies for deploying DL models: the two-step (2S) and the one-step (1S) approaches. The 2S approach consists of a classification stage to identify the active fingers and a regression stage to predict the trajectories of digits in motion. The 1S approach only has one regression stage to predict the trajectories of all fingers concurrently. In practice, the 2S approach should be marginal more efficient because not all models are inferred at a given moment. The models in the 2S approach are only trained on the subset where a particular is active, while all models in the 1S approach are trained on the full dataset. Here we focus on exploring the trade-offs between two approaches to inform future decisions of implementing the DL-based motor decoder in real-world applications.

The rest of this paper is organized as follows: section “Data Description” introduces the human participant of this research, the process of collecting input neural signals from the residual peripheral nerves of the participant, and establishing the ground-truth for the motor decoding paradigm using DL models. Section “Data Preprocessing” elaborates on how to cut raw input neural data into trials and extract their main features in the temporal domain before feeding to DL decoding models. Section “Proposed Methods and Deep Learning Models for Motor Intent Decoding” discusses the two approaches to efficiently translate motor intent from the residual peripheral nerves of the participant into motor control of the prosthesis as well as the architecture and the hyper-parameters of the DL models used in each approach. Section “Experimental Setup” is about the three ML models used as the baseline and how to input neural data are allocated to the training and validation set. Section “Metrics and Results” presents the metrics to measure the performance of all models used in the motor intent decoding process and discusses the main results of both proposed approaches. Section “Discussion” discusses the role of feature extraction in reducing the DL motor decoders’ complexity for real-time applications, how to further apply it in future works, and the advantages of ML and DL motor decoders in different scenarios where input dataset’s size varies. Finally, section “Conclusion” is to summarize the main contributions of this paper.

## 2 Data Description

### 2.1 Human Participant

The participant is a transradial male amputee who has lost his hand for over five years (Fig. 1). Among seven levels of upper-limb amputation, transradial is the most common type that accounts for about 57% of upper-limb loss in the U.S.(Schultz, 2011; Cordella, 2016). Like most amputees, the subject still has phantom limb movements; however, such phantom feelings fade away over time. By successfully decoding neural signals from the residual nerves of an amputee who has lost his limb for a long time, we would offer a chance to regain upper-limb motor control for those who are sharing the same conditions.

**Figure 1:**
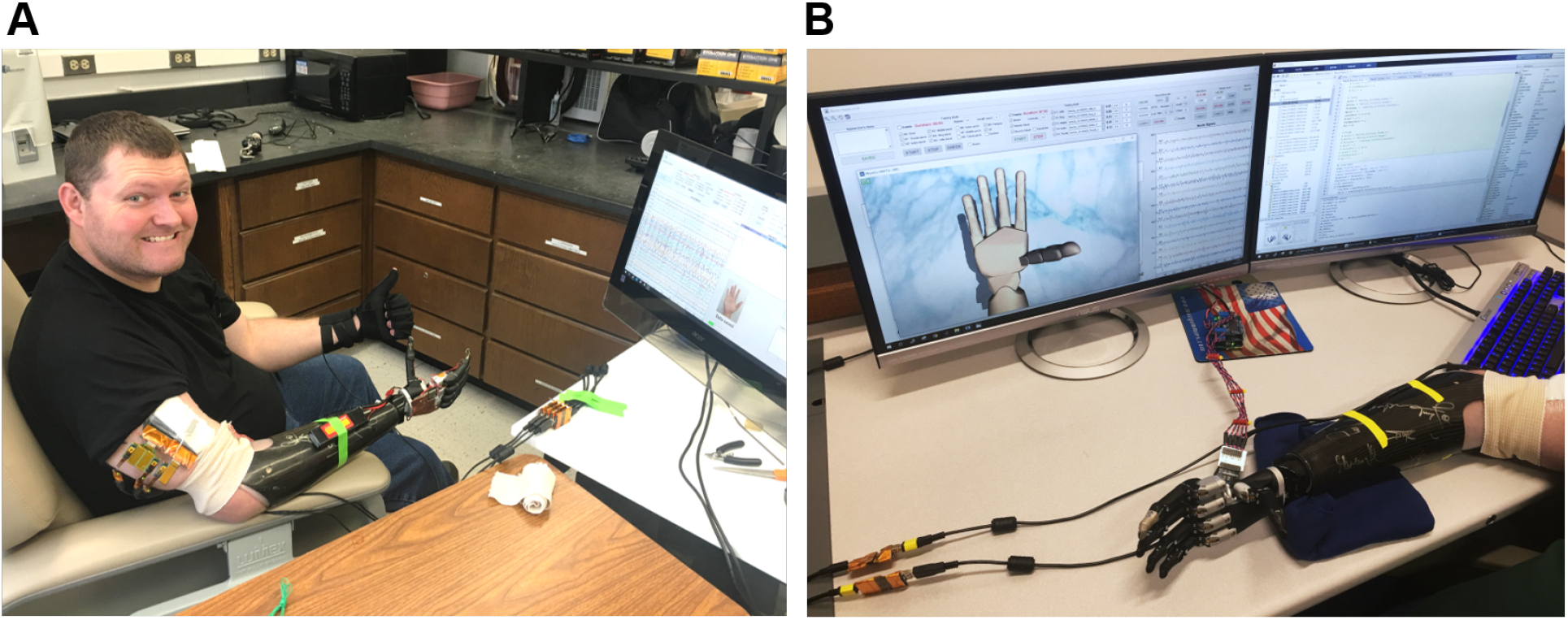
Photo of the (A) amputee and (B) the data collection software during a training session. The patient performs various hand movements repeatedly during the training session. Nerve data and ground-truth movements are collected by a computer and displayed in real-time on the monitor for comparison.

The human experiment protocols are reviewed and approved by the Institutional Review Board (IRB) at the University of Minnesota (UMN) and the University of Texas Southwestern Medical Center (UTSW). The patient voluntarily participates in our study and is informed of the methods, aims, benefits, and potential risks of the experiments before signing the Informed Consent. Patient safety and data privacy are overseen by the Data and Safety Monitoring Committee (DSMC) at UTSW. The implantation, initial testing, and post-operative care are performed at UTSW by Dr. Cheng and Dr. Keefer, while motor decoding experiments are performed at UMN by Dr. Yang’s lab. The clinical team travels with the patient in each experiment session.

The patient undergoes an implant surgery where four longitudinal intrafascicular electrode (LIFE) arrays are inserted into the residual median and ulnar nerves using the microsurgical fascicular targeting (FAST) technique (Fig. 2(A)). The electrode array’ design, characteristics, and surgical procedures are reported in (Cheng, 2017; Overstreet, 2019). The patient has the electrode arrays implanted for 12 months, during which the conditions of the implantation site is regularly monitored for signs of degradation.

**Figure 2:**
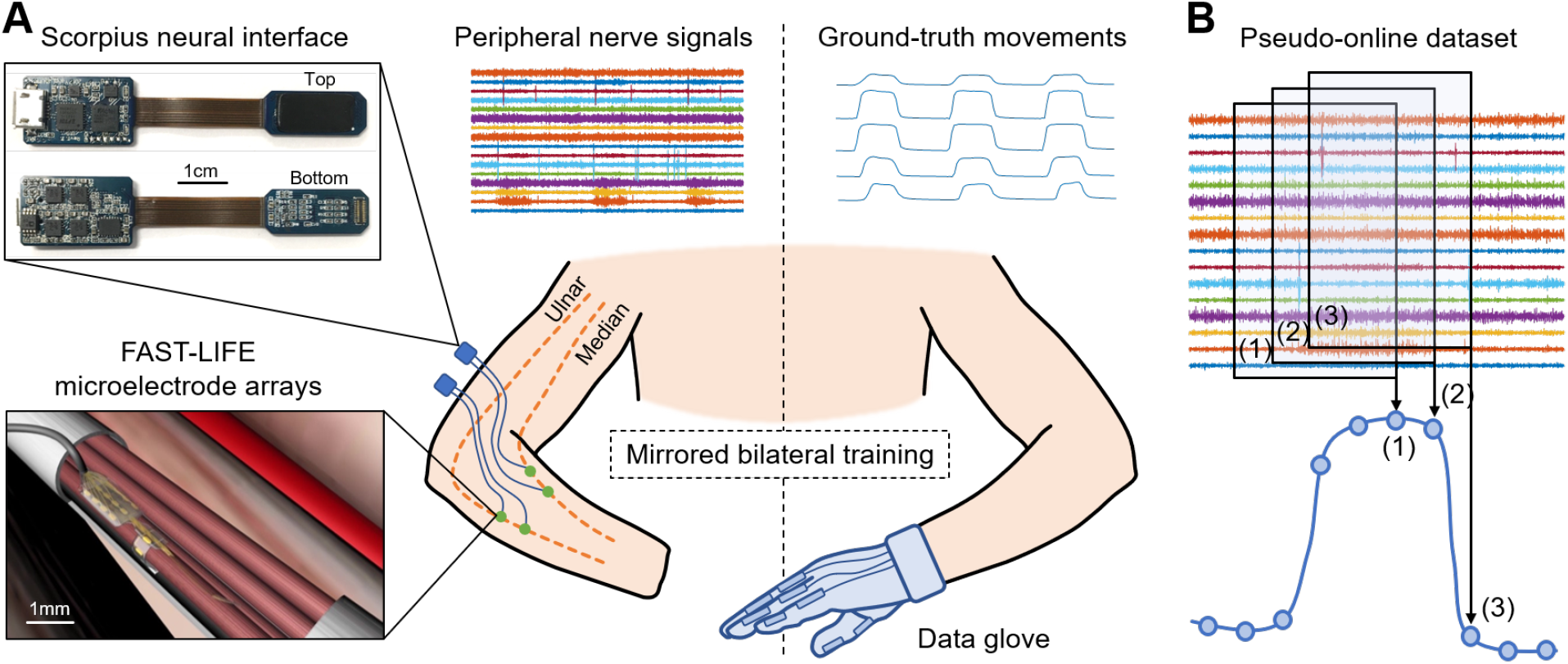
(A) Overview of the human experiment setup and data acquisition using the mirrored bilateral training. The patient has four FAST-LIFE microelectrode arrays implanted in the residual ulnar and median nerve (Overstreet, 2019). Peripheral nerve signals are acquired by two Scorpius neural interface devices (Nguyen & Xu, 2020). The ground-truth movements are obtained with a data glove. (B) Neural data are cut using a sliding window to resemble online decoding.

The patient participates in several neural stimulations, neural recording, and motor decoding experiment sessions. He initially has weak phantom limb movements due to reduced motor control signals in the residual nerves throughout the years. However, the patient reports that the more experiment sessions he takes part in, the stronger his phantom control and sensation of the lost hand become. This suggests that training may help re-establish the connection between the motor cortex and the residual nerves, resulting in better motor control signals.

### 2.2 Nerve Data Acquisition

Nerve signals are acquired using the Scorpius neural interface (Fig. 2(A)) - a miniaturized, high-performance neural recording system developed by Yang’s lab at UMN. The system employs the Neuronix chip family, which consists of fully-integrated neural recorders designed based on the frequency shaping (FS) architecture (Xu, 2014; Yang, 2016, 2018, 2020; Xu, 2020). The specifications of the Scorpius system are reported in (Nguyen & Xu, 2020). The system allows acquiring nerve signals with high-fidelity while suppressing artifacts and interference. Here two Scorpius devices are used to acquire signals from 16 channels across four microelectrode arrays at a sampling rate of 40 kHz (7.68 Mbps, 480 kbps per channel). The data are further downsampled to 5 kHz before applying a bandpass filter in 25-600 Hz bandwidth to capture most of the signals’ power. This results in a pre-processed data stream of 1.28 Mbps (80 kbps per channel).

### 2.3 Ground-truth Collection

The mirrored bilateral training paradigm (Sebelius, 2005; Jiang, 2012) is used to establish the ground-truth labels needed for supervised learning (Fig. 2(A)). The patient performs various hand gestures with the able hand while simultaneously imagining doing the same movement with the phantom/injured hand. The gestures include bending the thumb, index, middle, ring, little finger, index pinch, tripod pinch, and grasp/fist. Each gesture is repeated 100 times, altering between resting and flexing. Peripheral nerve signals are acquired from the injured hand with the Scorpius system, while ground-truth movements are captured with a data glove (VMG, 30, Virtual Motion Labs, TX) from the able hand. The glove can acquire up to 15 DOF; however, we only focus on the main 10 DOF (MCP and PIP) corresponding to the flexion/extension of five fingers.

## 3 Data Preprocessing

### 3.1 Cutting Raw Neural Data

In this paper, raw neural data are cut using a sliding window to resemble online motor decoding (Fig. 2(B)). Here the window’s length is set to 4 sec with an incremental step of 100 msec. At any instant of time, the decoder can only observe the past neural data. The pseudo-online dataset contains overlapping windows from a total of 50.7 min worth of neural recordings. Each of these 4-sec neural data segments serves as an input trial of the motor decoding process later.

### 3.2 Feature Extraction

Previous studies have shown that feature extraction is an effective gateway to achieve optimal classification performance with signals in the low-frequency band by highlighting critical hidden information while rejecting unwanted noise and interference. Here we select fourteen of the most simple and robust features that are frequently used in previous motor decoding studies (Zardoshti-Kermani, 1995; Phinyomark, 2009; Rafiee, 2011; Phinyomark, 2012). They are chosen such that there is no linear relationship between any pair of features. All features can be computed in the temporal domain with relatively simple arithmetic, thus aiding the implementation in future portable systems.

Table 1 summaries the descriptions and formula of the features. *x_i_* is the 4-sec neural data segments, which are further divided into windows of 100 milliseconds with *N* is the window length. Two consecutive windows are 80% overlapped, which is equivalent to a 20 milliseconds time step. This results in a data stream of 224 features over 16 channels with a data rate of 179.2 kbps (11.2 kbps per channel), which is more than 40 times lower than the raw data rate. Fig. 3 presents an example of the feature data in one trial that shows a clear correlation between the changes of the 14 extracted features and the finger’s movement. The amplitude of each feature is normalized by a fixed value before feeding to the DL models.

**Table 1:**
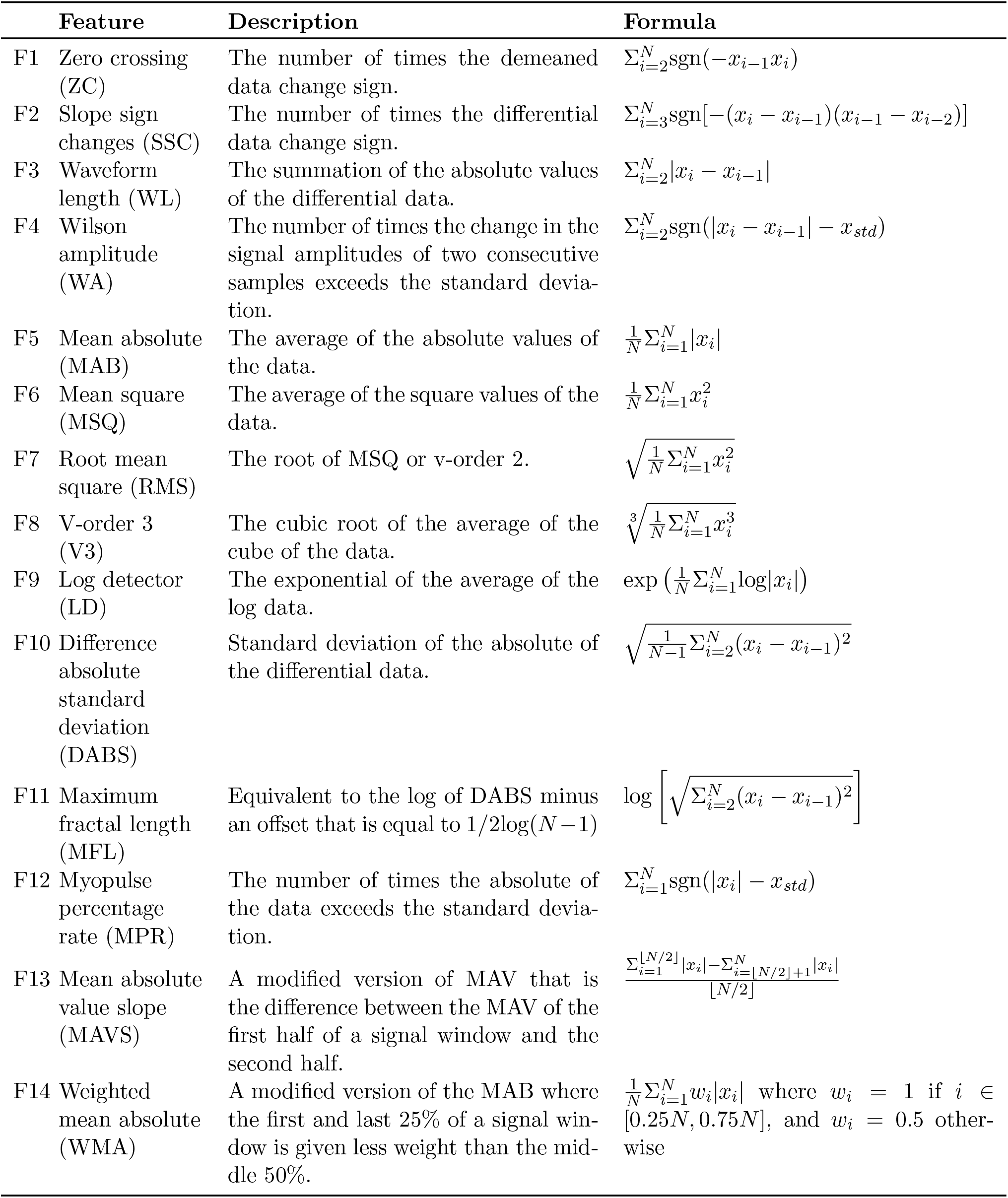
List of features, descriptions, and formula

**Figure 3:**
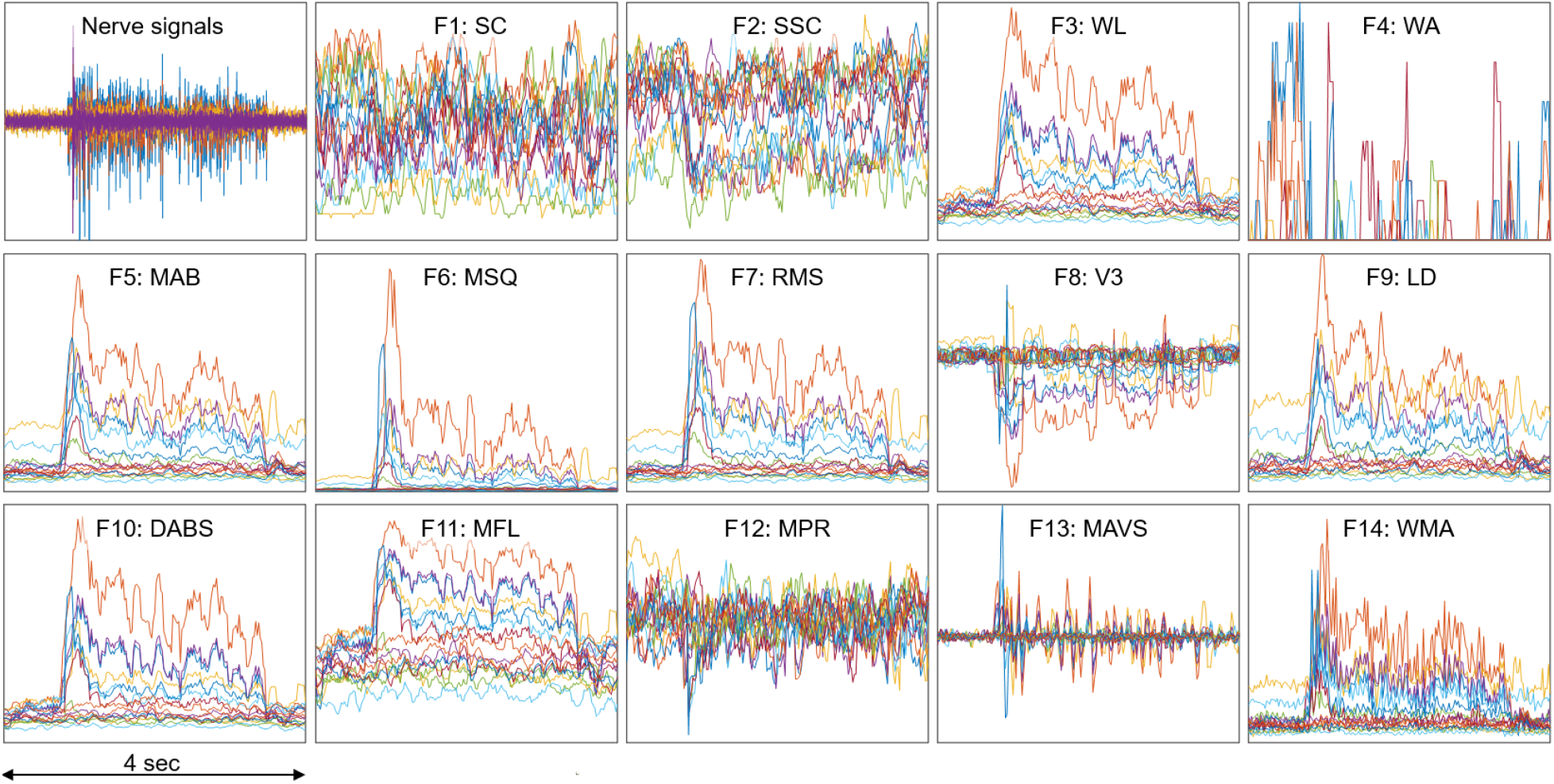
An example of feature data in one trial which show clear correlation with the finger’s movement. The amplitude of each feature is normalized by a fixed value.

## 4 Proposed Methods and Deep Learning Models For Motor Intent Decoding

### 4.1 Two-Step (2S) Strategy

Each finger exists in a binary state: active or inactive, depending on the patient’s intent to move it or not. There are 32 different combinations of five fingers corresponding to 32 hand gestures. Only a few gestures are frequently used in daily living activities, such as bending a finger (“10000”, “01000”,…, “00001”), index pinching (“00011”), or grasp/fist (“11111”). Therefore, classifying the hand gesture before regressing the fingers’ trajectories would significantly reduce the possible outcome and lead to more accurate predictions. The movement of inactive fingers could also be set to zero, which lessens the false positives when a finger “wiggles” while it is not supposed to.

Fig. 4(A) shows an illustration of the 2S strategy. In the first step, the classification output is a [1×5] vector encoding the state of five fingers. While the dataset used in this study only includes nine possible outcomes, the system can be easily expanded in the future to cover more hand gestures by appending the dataset and fine-tuning the models. In contrast, many past studies focus on classifying a specific motion, which requires modifying the architecture and re-training the models to account for additional gestures.

**Figure 4:**
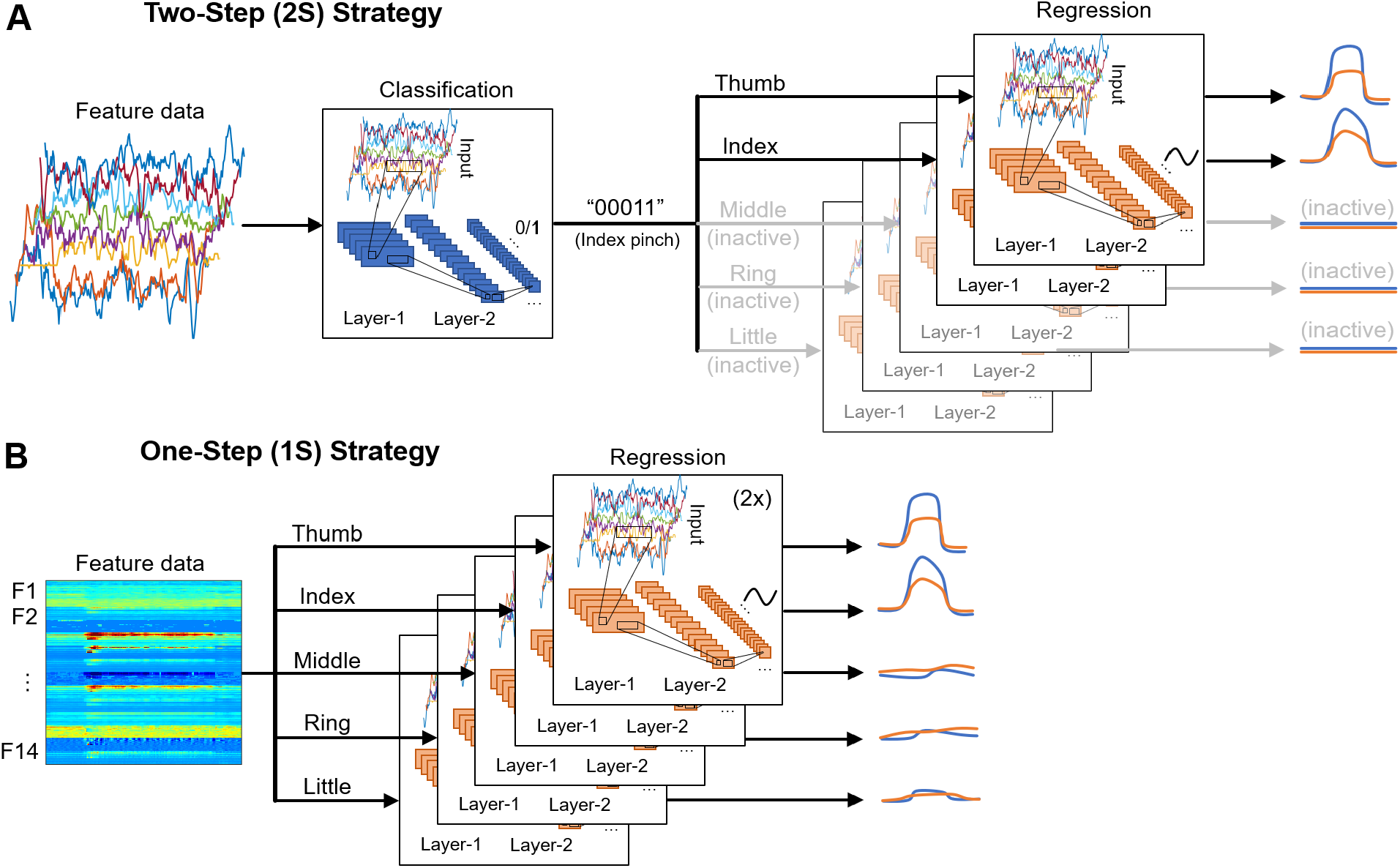
Illustration of the (A) two-step (2S) and (B) one-step (1S) strategy for deploying DL models.]

In the second step, the trajectory of each DOF is regressed by a DL model. Ten separate models regress the trajectory of ten DOF (two per finger). The models associated with inactive fingers are disabled, and the prediction outputs are set to zero. As a result, the dataset used to train each model is only a subset of the full dataset where the corresponding DOF is active. While all models use the same architecture, they are independently optimized using different sets of training parameters such as learning rate, minibatch size, number of epochs, etc., to achieve the best performance. An advantage of this approach is that if one DOF fails or has poor performance, it would not affect the performance of others.

### 4.2 One-Step (1S) Strategy

Fig. 4(B) shows an illustration of the 1S strategy. It is the most straightforward approach where the trajectories of each DOF are directly regressed regardless of the fingers’ state. As a result, the full dataset, which includes data when the DOF is active (positive samples) and idle (negative samples), must be used to train each DOF. Because the number of negative samples often exceeds the number of positive samples from 5:1 to 10:1, additional steps such as data augmentation and/or weight balancing need to be done during training. This also leads to more false-positives where an idle DOF still has small movements that could affect the overall accuracy.

Moreover, it is worth noting that the time-latency and efficiency of the 1S approach do not necessarily better than the 2S. In the second step of the 2S approach, several DL models are disabled depending on the hand gesture, resulting in lower overall latency and computation in most implementations where there is only one processing unit (GPU or CPU). The actual performance differences are difficult to be quantitatively measured because the proportion of individual hand gestures largely depends on the user and scenarios. Nevertheless, by comparing the decoding outcomes of the 1S and 2S strategy, we can better understand the trade-off between simplicity and accuracy.

### 4.3 Deep Learning Models

Fig. 5 shows the architecture of the DL classification and regression models. They include standard building blocks such as convolutional, long-short term memory (LSTM), fully-connected, and dropout layers of different combinations, order, and set of parameter values. The architecture is optimized by gradually adding layers and tuning their parameters while tracking the decoder’s efficacy using 5-fold cross-validation. As the performance converges, additional layers would tend to result in over-fitting.

**Figure 5:**
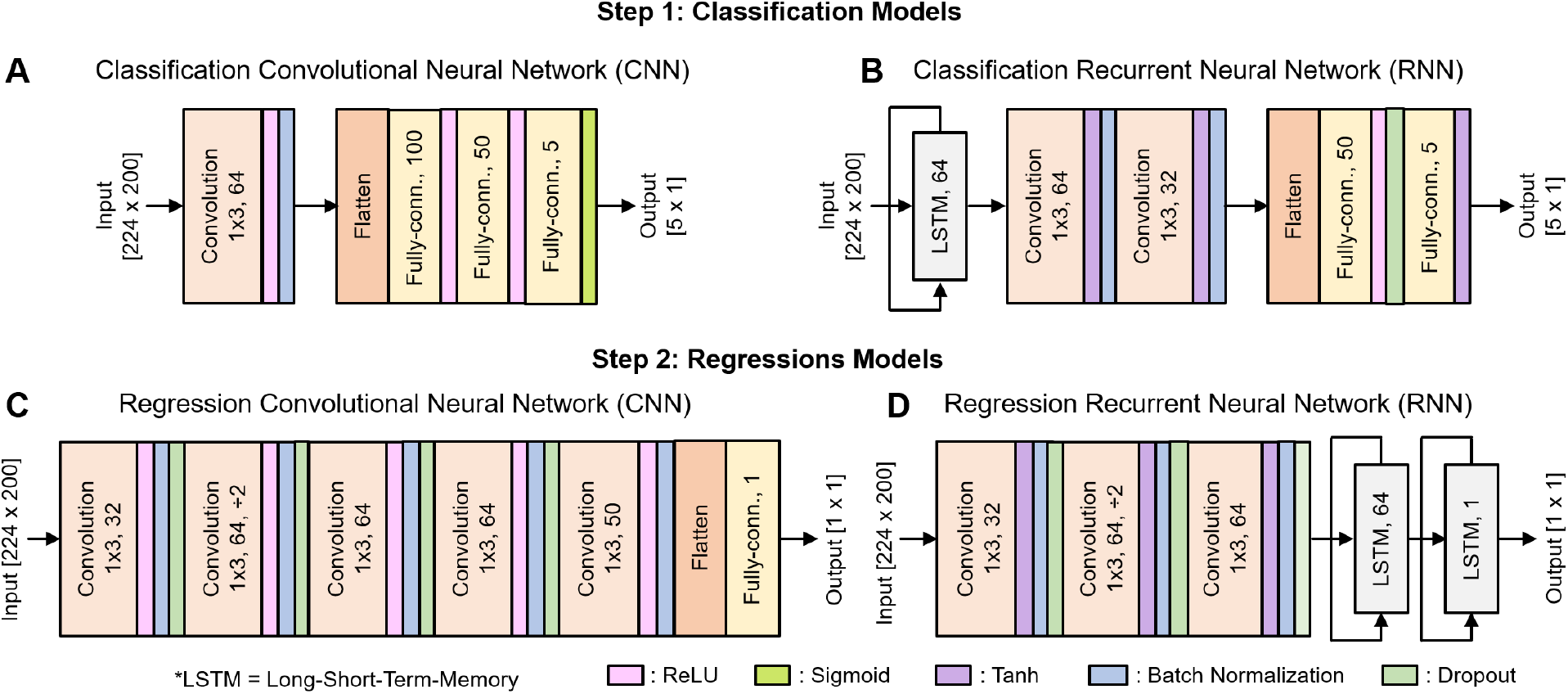
Architecture of the DL models: (A) CNN for classification, (B) RNN for classification, (C) CNN for regression, (D) RNN for regression.

There are ten copies of the regression model for ten DOF, each of which is trained separately. We use Adam optimizer with the default parameters *β*_1_ = 0.99, *β*_2_ = 0.999, and a weight decay regularization *L*_2_ = 10^−5^. The mini-batch size is set to 38, with each training epoch consists of 10 mini-batches. The learning rate is initialized to 0.005 and reduced by a factor of 10 when the training loss stopped improving for two consecutive epochs.

Table 2 shows a rough comparison between the DL models used in this and our previous work. Note that for regression, the number of learnable parameters is a total of ten models for ten different DOF. The addition of feature extraction, thus dimensional reduction, allows significantly lowing the DL models’ size and complexity. This is essential for translating the proposed decoding paradigm into a real-time implementation for portable systems.

**Table 2:** Comparison between this work and Nguyen & Xu *et al.* (2020)

|  | No. of layers |  |  | No. of parameters |
| --- | --- | --- | --- | --- |
|  | Conv. | LSTM | Fully conn. |  |
| Nguyen & Xu *et al.* (2020) | 21 | 2 | 3 | 25,927,050 |
| This work (classification) | 1 | 0 | 3 | 1,465,749 |
| This work (regression) | 3 | 2 | 0 | 767,200 |

## 5 Experimental Setup

In this research, we investigate the performance of two main DL architectures: the CNN and RNN for both classification and regression tasks. Besides, the DL models are benchmarked against “classic” supervised learning techniques as the baseline. They include support vector machine (SVM), random forest (RF), and multi-layer perceptron (MLP).

For baseline techniques, the input of the classification task is the average of 224 features across 200 time-steps, while the input of the regression task is the 30 most important PCA components. The SVM models use the radial basis function (RBF) for classification and polynomial kernel of degree three for regression, with parameter *C* = 1. The RF models use five and ten trees for classification and regression, respectively, with a max depth of three. The MLP model for classification is created by replacing the convolutional layer of the CNN model with a fully-connected layer of 200 units. The MLP model for regression has four layers with 300, 300, 300, and 50 units, respectively.

The 5-fold cross-validation is used to compare the performance of the classification task. For the regression task, the dataset is randomly split with 80% for training and 20% for validation. The split is done such that no data windows from the training set to overlap with any data windows from the validation set.

## 6 Metrics and Results

### 6.1 Metrics

This subsection introduces the metrics to measure the performance of five models, including the two discussed DL models and three other classic supervised learning techniques as benchmarks in both classification and regression tasks. The performance of the classification task is evaluated using standard metrics including accuracy and F1 score derived from true-positive (TP), true-negative (TN), false-positive (FP), and false-negative (FN) as follows:

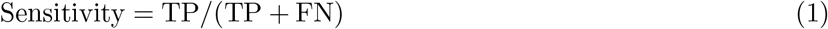

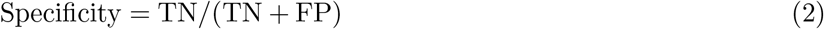

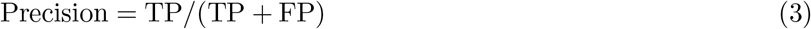

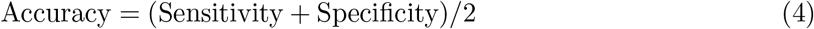

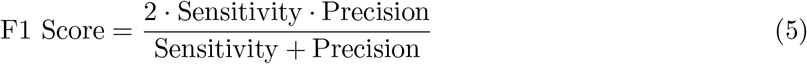

We use the definition of accuracy with an equal weight of sensitivity and specificity because the occurrence of class-1 (active finger) is largely outnumbered by the occurrence of class-0 (inactive finger).

The performance of the regression task is quantified by two metrics: mean squared error (MSE) and variance accounted for (VAF). MSE measures the absolute deviation of an estimated from the actual value of a DOF while VAF reflects relative deviation from the actual values of several DOF. They are defined as follows:

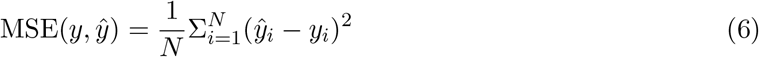

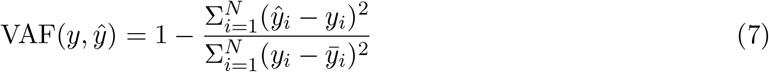

where *N* is the number of samples, *y* is the ground-truth trajectory, 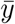 is the average of *y*, and 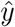 is the estimated trajectory. The value of *y* and 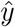 are normalized in a range [0, 1] in which 0 represents the resting position.

Although the MSE is the most common metric and effectively measures absolute prediction errors, it cannot reflect the relative importance of each DOF to the general movements. For example, if the average magnitude of DOF A is hundreds of times smaller than that of DOF B, a bad estimation of A still can yield lower MSE than a reasonable estimation of B. The VAF score is more robust in such scenarios; thus, it could be used to compare the performance between DOF of different magnitude. The value of the VAF score ranges from (−*∞*, 1], the higher, the better. For practicality, negative VAF values are ignored when presenting the data.

### 6.2 Classification Results

Table 3 shows the average 5-fold cross-validation classification results of all the techniques. The predictability of each finger is largely different from one another. The thumb, which produces strong signals only on the median nerve (first eight channels), is easily recognized by all techniques. Overall, CNN offers the best performance with accuracy and an F1 score for all finger exceeding 99%. RF closely follows with performance ranging from 98% to 99%. While DL still outperforms classic techniques, it is worth noting that RF could also be a prominent candidate for real-world implementation because RF can be more efficiently deployed in low-power, portable systems.

**Table 3:**
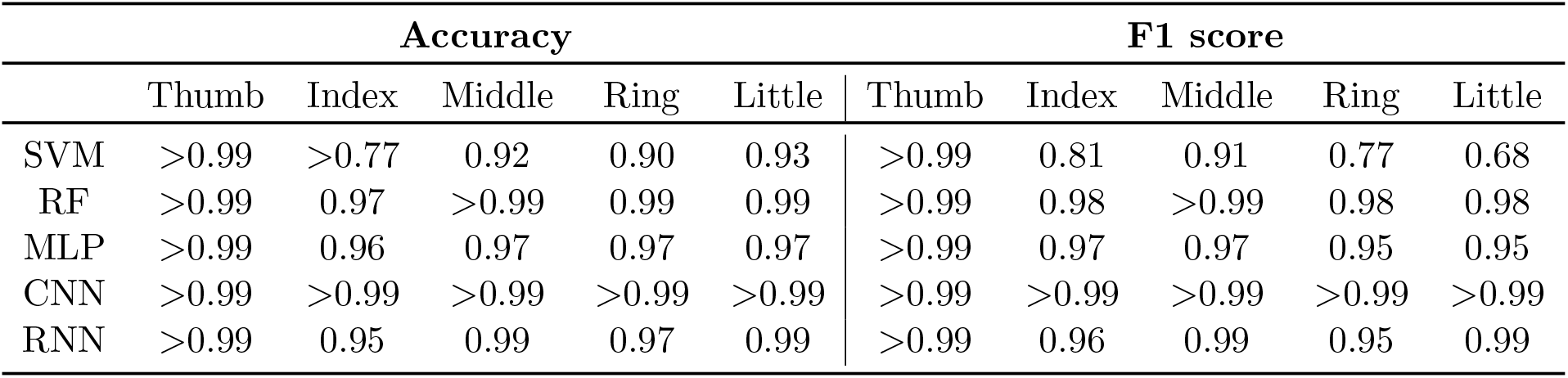
Classification performance

### 6.3 Regression Results

Fig. 6 presents the MSE and VAF scores for both strategies. In the 1S approach, the DL models, especially RNN, significantly outperform the other methods in both MSE and VAF, as shown in Fig. 6(A, C). In the 2S approach, the performance is more consistent across all methods, where classic methods even outperform DL counterparts in certain DOF as shown in Fig. 6(B, D). Between the two strategies, the 1S approach generally gives better results; however, the high performance can only be achieved with RNN.

**Figure 6:**
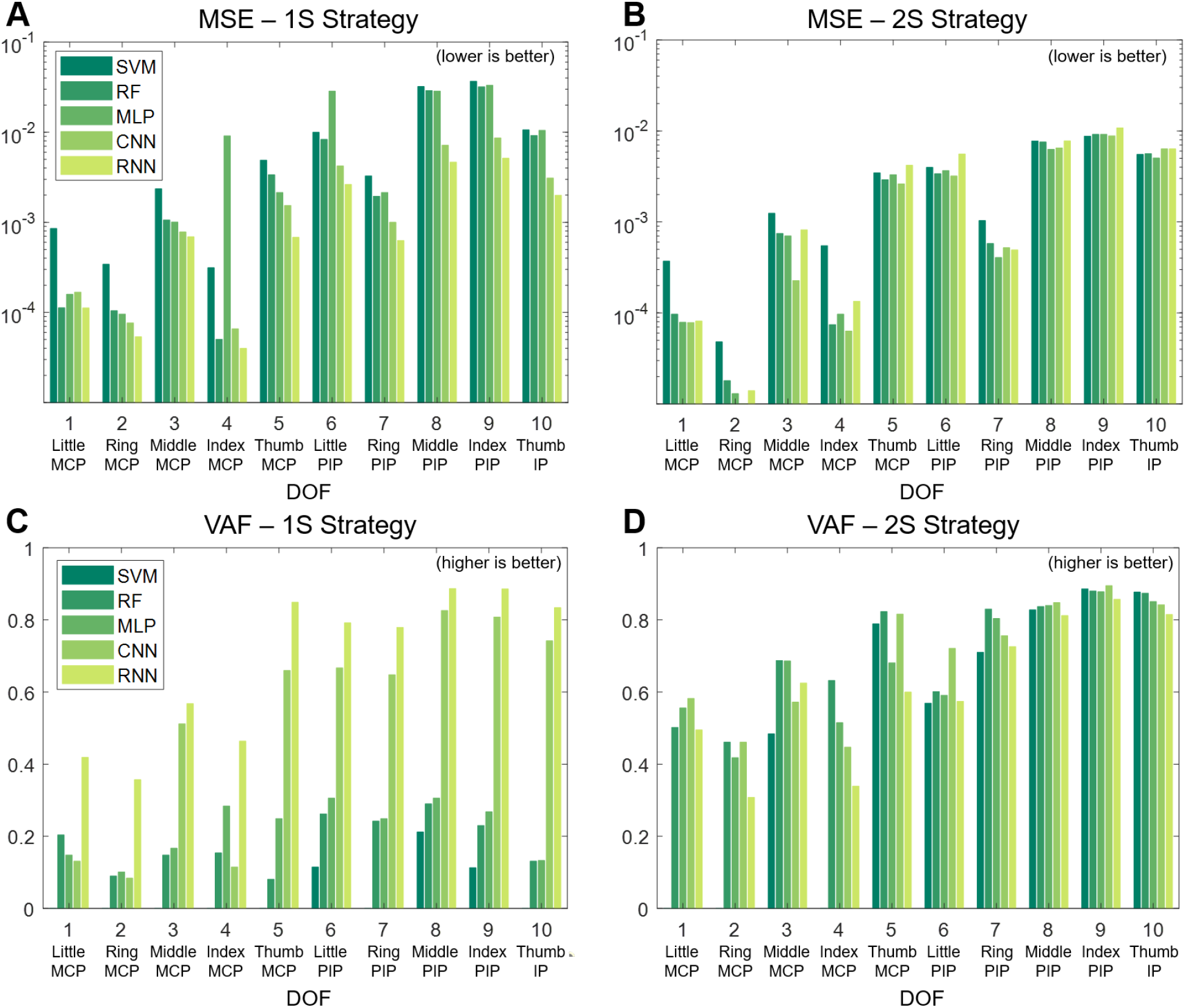
Regression performance in term of MSE (A, B) and VAF (C, D).

## 7 Discussion

### 7.1 Feature Extraction Reduces Decoders’ Complexity

Both DL architectures investigated in this study, namely CNN and RNN, delivers comparable motor decoding performance to our previous work (Nguyen & Xu, 2020) while require much lower computational resources to implement. The average VAF score for most DOF is 0.7, with some exceeding 0.9. Such results are achieved with relatively shallow DL models with 4-5 layers, a significant reduction from the previous implementation with 26 layers. While an extra step of feature extraction is required, all feature extraction techniques are specifically designed to be efficiently computed with conventional arithmetic processors (e.g., CPU) and/or hardware accelerators (e.g., FPGA, ASIC).

Another direction for future works would be including additional features. Here we apply the most common features in the temporal domain for their simplicity and well-established standing for neuroprosthesis applications. There are other features such as mean power, median frequency, peak frequency, etc. in the frequency domain that have been explored in previous literature. The purpose of this study is not to exhaustively investigate the effect of all existing features in motor decoding but to prove the effectiveness of the feature extraction method in achieving decent decoding outcomes. The success of this method will open up to future research on simultaneously applying more features in multiple domains for better outcomes.

Furthermore, it is worth noting that the use of feature extraction excludes most of the high-frequency band 600-3000 Hz, which is shown in our previous work that could contain additional nerve information associated with neural spikes. A future direction would be extracting that information using spike detection and sorting technique and combine them with the information of the low-frequency band to boost the prediction accuracy. However, the computational complexity must be carefully catered to not to hinder the real-time aspect of the overall system.

### 7.2 Classic Machine Learning v.s. Deep Learning

The classification task can be accomplished with high accuracy using most classic ML techniques (e.g., RF) and near-perfect with DL approaches (e.g., CNN). Along with other evidence in (Nguyen & Xu, 2020), this suggests that nerve data captured by our neural interface contain apparent neural patterns that can be clearly recognized to control neuroprostheses. While the current dataset only covers 9/32 different hand gestures, these are still promising results that would support future developments, including expanding the dataset to cover additional gestures.

The regression results are consistent with the conclusion of many past studies that DL techniques only show clear advantages over classic ML methods when handling a large dataset. This is evident in the 1S strategy, where each DOF is trained with the full dataset consisting of all possible hand gestures. In contrast, in the 2S strategy, where the dataset is divided into smaller subsets, DL techniques lose their leverage. However, as the dataset is expanded in the future, we generally believe that DL techniques should emerge as the dominant approach.

## 8 Conclusion

This work presents several approaches to optimize the motor decoding paradigm that interprets the motor intent embedded in the peripheral nerve signals for controlling the prosthetic hand. The use of feature extraction largely reduces the data dimensionality while retaining essential neural information in the low-frequency band. This allows achieving similar decoding performance with DL architectures of much lower computational complexity. Two different strategies for deploying DL models, namely 2S and 1S, with a classification and a regression stage, are also investigated. The results indicate that CNN and RF can deliver high accuracy classification performance, while RNN gives better regression performance when trained on the full dataset with the 1S approach. The findings layout an important foundation for the next development, which is translating the proposed motor decoding paradigm to real-time applications, which requires not only accuracy but also efficiency.

## Acknowledgments

The surgery and patients related costs were supported in part by the Defense Advanced Research Projects Agency (DARPA) under Grants HR0011-17-2-0060 and N66001-15-C-4016. The technology development and human experiments were supported in part by internal funding from the University of Minnesota, in part by the NIH under Grant R21-NS111214, in part by NSF CAREER Award No. 1845709, and in part by Fasikl Incorporated.

Zhi Yang is co-founder of, and holds equity in, Fasikl Inc., a sponsor of this project. This interest has been reviewed and managed by the University of Minnesota in accordance with its Conflict of Interest policy. J. Cheng and E. W. Keefer have ownership in Nerves Incorporated, a sponsor of this project.

